# Process optimization for enhanced enzymatic nylon deconstruction

**DOI:** 10.1101/2025.11.26.690883

**Authors:** Célestin Bourgery, Lance X. Zhang, Dana L. Carper, Muchu Zhou, Robert L. Sacci, Matt Korey, Lei Wang, Jessica H.B. Hemond, John F. Cahill, Vera Bocharova, Joshua K. Michener

## Abstract

Plastics such as polyamides (PAs) possess unique physicochemical properties that make them indispensable in modern society. However, their energy-intensive production and challenging end-of-life management highlight the urgent need for efficient recycling or remanufacturing solutions. Enzymatic depolymerization offers a promising route toward circular recycling, yet remains constrained by limited enzyme characterization, lack of validation under industrially relevant conditions, and overall performance. Here we optimized the reaction conditions for three recently discovered nylon-degrading enzymes. One of them, Nyl12, achieved product titers with PA6 and PA66 that exceed previously reported values, without enzyme engineering or substrate pretreatment. We further demonstrated the scalability of the process and its application to complex PA-based materials used in microelectronic components. Analysis of substrate features, including surface area and particle size, revealed key parameters governing enzymatic activity and provided a framework for future pretreatment and process optimization efforts. In combination, these efforts provide a new benchmark for enzymatic nylon recycling.

## Introduction

Plastics combine exceptional manufacturability, tunable properties, and durability, making them indispensable to modern society. Their low density, high specific stiffness, and resistance to corrosion and chemicals provide an unmatched performance-to-cost ratio, features that explain their ubiquity and underscore the urgency of developing efficient remanufacturing strategies rather than merely reducing their use.^[1–4]^

However, the recycling of plastics has become a critical component of contemporary waste management and sustainability initiatives, driven by growing concerns over waste accumulation and the substantial energy consumption and greenhouse gas emissions associated with plastic production. Moreover, accumulated plastic waste has potential as a feedstock for (re)manufacturing, such as the production of new value-added products or novel materials. ^[3–5]^

Nylons, particularly polyamide 6 (PA6) and polyamide 6,6 (PA66), are among the most widely used synthetic polymers. They play pivotal roles in industries such as textiles, automotive, and electronics. However, despite their extensive use, Nylon recycling is challenging, largely due to the strength of the amide bond, the high crystallinity of the polymer, and the presence of additives and contaminants. In combination, these factors make Nylons resistant to conventional recycling methods.

In recent years, enzymatic plastic recycling has emerged as a promising alternative approach. Work on polyethylene terephthalate (PET), including the discovery and engineering of novel enzymes combined with process optimization, has enabled efficient degradation and recycling of this polymer.^[6–13]^ While these efforts have inspired the exploration of new systems for recycling other polymers, enzymatic hydrolysis of Nylons remains limited. Beginning in the 1980s, Negoro and colleagues discovered and characterized the first nylon oligomer hydrolases, NylA (a 6-aminohexanoate cyclic dimer hydrolase), NylB (a 6-aminohexanoate oligomer hydrolase), and NylC (a nylon oligomer endohydrolase).^[14–17]^ Collectively, these enzymes demonstrated the ability to hydrolyze PA6 oligomers into shorter chains and monomers, providing the first biochemical evidence that synthetic polyamides could be enzymatically degraded. More recent studies have advanced the understanding of the substrate and product specificity of novel nylon hydrolases, revealing their potential for more efficient recycling processes.^[18–22]^

Recently, we identified new NylC homologs capable of degrading PA6 and PA66.^[19]^ Three new enzymes, termed Nyl10, Nyl12, and Nyl50, were particularly promising. Their varied substrate- and product-selectivity offer the potential for targeted Nylon recycling. However, despite the considerable potential of nylon hydrolases, their application remains limited due to insufficient characterization and relatively low hydrolysis yields under the conditions tested. As suggested by numerous prior studies on enzymatic PET hydrolysis, increasing hydrolysis yields will require a combination of polymer pretreatment, process optimization, and enzyme engineering, aimed at altering polymer characteristics such as polymer crystallinity, molecular weight, surface area or dispersity to improve polymer accessibility and enzyme catalytic efficiency.^[13,23–39]^ Furthermore, in contrast to work on PET, enzymatic depolymerization of Nylons still suffers from a lack of studies using industrially-relevant conditions such as high substrate loadings, larger reaction scales, and more complex feedstocks.

In this study, we first optimized the reaction conditions for Nyl10, Nyl12, and Nyl50. The process was then successfully scaled up at process-relevant substrate loadings. We next examined key substrate properties, including surface area and particle sizes, to identify those that facilitate optimal enzymatic activity. The effect of surfactants and co-solvents were also assessed. Finally, we demonstrated a proof-of-concept for the enzymatic degradation of a mixed polyamide and PBT resin, including additives commonly used in microelectronics components. The characterization of optimal enzyme parameters and substrate properties provides a valuable foundation for engineering more efficient enzymes, which, when coupled with substrate pretreatment, could lead to more effective and scalable nylon recycling methods.

## Results and Discussion

### Characterization and determination of optimal reaction conditions for Nyl10, Nyl12, Nyl50

To assess the effects of process conditions on nylon hydrolase activity, we systematically varied the temperature (35°C to 85°C), pH (4 to 9), and buffer composition (phosphate *vs*. Tris) for the enzymatic hydrolysis of PA66 and PA6. Reactions were carried out with 10 mg of PA in 75 μL buffer plus 25 μl enzyme lysate, for 12 and 72 hours. Each enzymatic reaction and parameter were treated separately and compared with a reference setup (65°C, 20 mM phosphate buffer, pH 7.4).^[19]^

For all three enzymes, the titers of hydrolysis products increased with temperature, reaching an optimum at 75°C before sharply declining at 85°C, likely due to thermal inactivation of the enzyme (Figure 1A). This temperature profile underscores the high thermostability of the enzymes, enabling hydrolysis reactions to be performed at or above the glass transition temperature (T_g_) of the polyamides and thereby enhancing polymer accessibility and degradation efficiency (Figures 1A, S1A and S1B).^[32,39,40]^ The enzymes exhibited hydrolytic activity across the entire tested pH range (pH 4 to 9), with optimal performance observed between pH 6 and 9, peaking at pH 8 (Figure 2B). Under equivalent pH conditions, the use of 200 mM Tris buffer resulted in a 2-fold increase in hydrolysis titer at pH 7 and a 6-fold increase at pH 8 compared to a 200 mM phosphate buffer (Figure S1C). Minimal non-enzymatic hydrolysis was detected in our control. Similarly, the addition of 10 mM metal ions (MgCl_2_, CaCl_2_, ZnSO_4_, or NiSO_4_) did not result in any increase in product titers, while certain ions, such as ZnSO_4_ and NiSO_4_, even led to a reduction in titers (Figure S2). We did not evaluate the impact of additional buffer alternatives in this study. However, if needed, a broader screening could be conducted in the future to better tailor the reaction conditions to downstream isolation and purification requirements for the hydrolysis products.

**Figure 1.**
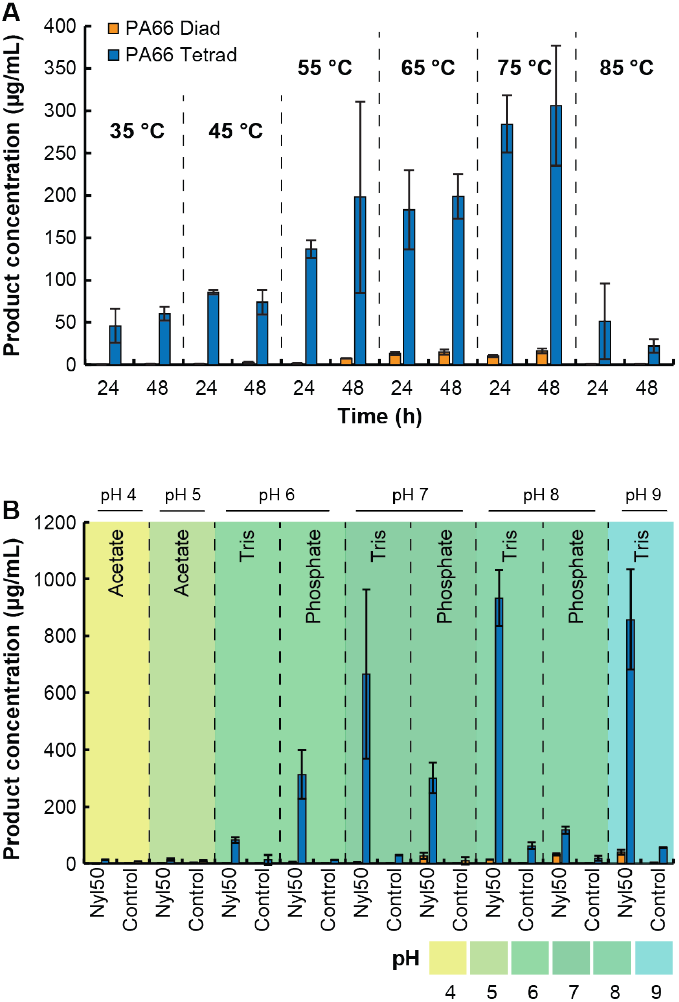
Enzymatic PA66 depolymerization reactions with Nyl50 in function of temperature and pH. (A) Effect of temperature on Nyl50 hydrolysis activity. Reactions ran with crude lysate in 20 mM phosphate pH 7.4 for 24 h. (B) Effect of pH on hydrolysis activity. Reactions ran with crude lysate for 24 h at 65°C. In both panels, error bars show the standard deviation, calculated from three biological replicates.

**Figure 2.**
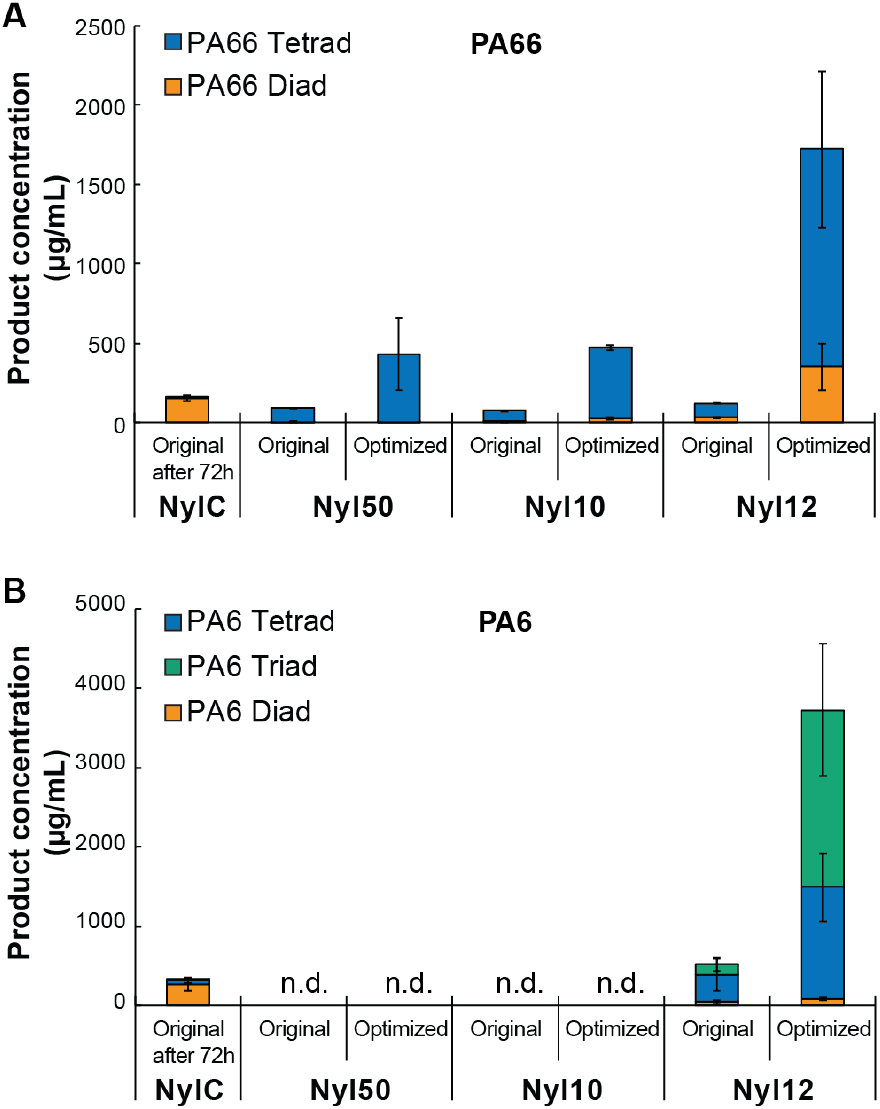
Enzymatic PA66 and PA6 depolymerization reactions with Nyl50, Nyl10 and Nyl12 under optimized conditions. (A) PA66 (B) PA6. Reactions were conducted with 0.3 mg/mL of pure enzyme, “Original” in 20 mM Phosphate pH 7.4 at 65°C and “Optimized” in 200 mM Tris pH 8 at 75°C, both for 24 h unless otherwise specified. Here, NylC is the quadruple mutant NylC-GYAQ. Error bars show the standard deviation, calculated from three biological replicates. n.d.: not detected.

### Partial PAs Degradation Under Optimal Conditions Establishes Nyl12 as the New Benchmark

The optimal reaction conditions identified in this study (75°C, pH 8, Tris buffer) were then implemented in combination and compared to the original conditions (65°C, pH 7.4, phosphate buffer). Reactions were conducted using a consistent pure enzyme concentration of 0.3 mg/mL and 10 mg of washed PA66 or PA6 powder in a total volume of 100 µL, in alignment with prior work.^[19]^

Under these optimized conditions, hydrolysis of PA66 using Nyl50 resulted in titers of approximately 500 mg/L after both 24 and 72 h, while titers for Nyl10 reached approximately 700 mg/L after 72 h. These values represent 4.7-fold and 3.2-fold increases for Nyl50 at 24 and 72 h, respectively, and 6.4-fold and 2.7-fold increases for Nyl10, relative to the previous conditions, while no activity was observed with PA6 (Figure 2 and S3). Similarly, for PA6 using Nyl12, hydrolysis titers of approximately 4000 mg/L were observed after both 24 and 72 h, corresponding to >7-fold improvement after 24 h and a 3.8-fold improvement after 72 h compared to the reference condition. These results correspond to degrees of depolymerization of approximately 0.7% for PA66 and 4% for PA6. For PA66, these results are comparable to the highest reported in the current literature using engineered enzymes,^[18,41]^ while the degree of depolymerization of PA6 with Nyl12 outperforms previous studies. Furthermore, a 24 h incubation of PA66 with Nyl12 resulted in approximately 2% degradation (1721 ± 637 µg/mL), positioning Nyl12 as the most efficient enzyme reported to date for both PA66 and PA6 (Figure 2). This establishes a new benchmark for enzymatic degradation in this substrate class. We also consider it important to specify that this enhanced depolymerization was achieved without the use of acid pretreatment of the polymer substrates, a step employed in previous studies as a pretreatment to increase enzyme accessibility.^[21,42]^

### Reaction scale up

We next considered scalability, using Nyl50 as a model system, as it is the best-characterized enzyme to date.^[19]^ To avoid enzyme purification, a step that increases complexity, time, and cost in large-scale applications, we sought to exploit the inherent thermostability of Nyl50 compared with the heat-labile native enzymes of its *E. coli* expression host. Crude lysates were subjected to thermal pretreatment at 60 °C or 70 °C for 10, 30, or 60 min to selectively precipitate host-derived components. Substantial precipitation was observed within 10 min at 60 °C, consistent with rapid denaturation of *E. coli* proteins, while the supernatant retained full hydrolytic activity relative to untreated controls (Figure S5A and S5B). Thermal pretreatment thus efficiently removed host-derived material without compromising Nyl50 activity and was adopted for all subsequent large-scale experiments.

Next, we optimized lysate concentration to maximize hydrolysis titer, evaluating the effect of lysate-to-buffer ratio using the thermally pretreated lysate. Reactions were carried out in a total volume of 100 µL while varying the lysate volume relative to the buffer. Increasing lysate volume enhanced hydrolysis titer, reaching an optimum at 25 µL lysate and 75 µL buffer. A 1:1 ratio with buffer (50 µL lysate and 50 µL buffer) led to an approximately 2-fold decrease in titer, likely due to reduced buffering capacity at higher lysate concentrations, thereby compromising the reaction environment (Figure S5C). Finally, we scaled up the reaction, with a constant substrate concentration, from 10 mg of PA66 in a 100 µL reaction volume to 100 mg in 1 mL under static conditions. Scale-up resulted in a very slight increased hydrolysis titer and less variability, likely due to improved contact between the solid polymer and the aqueous buffer phase. Further scale-up to 1 g PA66 in 10 mL reaction volume, conducted in a round-bottom flask under continuous agitation, yielded a comparable titer, demonstrating the robustness and scalability of the process (Figure S6A).

### Determination of substrate properties, surfactants and solvents addition, influencing enzymatic hydrolysis

To further enhance hydrolysis yield, the reactions were repeated using an increased enzyme concentration (1 mg/mL compared to 0.3 mg/mL). After 72 h, similar overall titers were observed for both enzyme systems. However, a shift in the oligomer distribution was detected, indicating progressive degradation of longer-chain oligomers into shorter fragments such as conversion of linear PA66 tetrads to PA66 diads or linear PA6 tetrads to PA6 triads and diads (Figure S7). These findings suggest that, under the tested conditions, the rate-limiting factor in the enzymatic hydrolysis of PA66 and PA6 is substrate accessibility rather than enzyme robustness or catalytic efficiency, consistent with recent observations reported by Bell et al. ^[18]^

Based on these observations, we chose to evaluate the effect of surface area and particle size of the substrate, which could provide critical clues for improving enzymatic degradation.^[27]^ Nyl50 with PA66 was selected as the model system for these investigations, extending our previous efforts using NylC.^[43]^ First, a range of PA66 samples with various surface areas were incubated with Nyl50 for 6 h under the optimal conditions identified above (75°C, pH 8, Tris buffer). The concentrations of both diads and tetrads increase with surface area up to 7.15 m^2^/g, beyond which the trend reverses and both concentrations decrease (Figure 3A). These results are consistent with the behavior of NylC in our previous study.^[43]^

**Figure 3.**
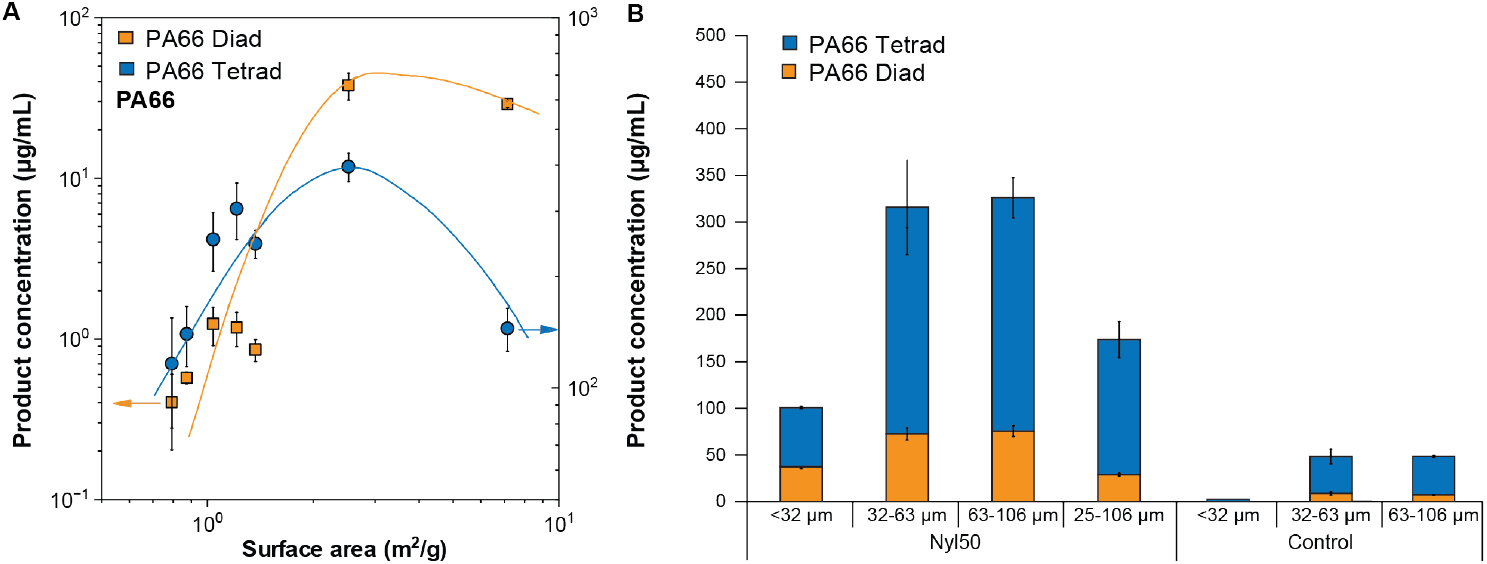
Determination of substrate properties influencing enzymatic hydrolysis. A) Concentration of hydrolysis products (diads and tetrads) of PA66 by Nyl50 plotted vs PA66 surface area. The dashed lines indicate the overall trend. B) Concentration of hydrolysis products *vs*. PA66 sizes in the presence and absence (control) of Nyl50. Reactions were performed with crude lysate, 6 h at 75°C. Error bars show the standard deviation, calculated from two biological replicates.

Generally, ball-milled PA66 powder contains a variety of particle sizes with a broad size distribution. To investigate the effect of particle size on enzyme activity, the ball-milled powder was sieved using specific cut-off sizes. Specifically, particles in the 25–106 µm size range were initially sieved from the powder, followed by further refinement through additional sieving, which produced narrower particle size fractions. The activity of Nyl50 was reduced using substrate particles <32 µm compared to size ranges 32–63 µm, 63–106 µm, and even broader composite particle range 25–106 µm (Figure 3B). Inhibition associated with smaller particle sizes may explain the observed decrease in activity with increasing surface area presented (Figure 3A). Similar trends were observed with Nyl12 (Figure S4) However, the observed effects could hypothetically be linked to a substrate-specific inhibitory mechanism: particles of this size remain dispersed longer than larger ones, which sediment rapidly. This potentially enhances substrate surface accessibility for enzymes, which also would increase the likelihood of non-specific and non-productive interactions between enzymes and substrates. Our results are consistent with the study conducted on PET by Brizendine *et al*., which demonstrated that particle size reduction increases the rate of enzymatic depolymerization but does not improve the overall conversion extent.^[27]^

We also investigated the potential of enzymes to adsorb onto PAs surfaces. Previous studies have shown that enzymes are often poorly adsorbed onto highly crystalline PET (crystallinity ≈48.6%) due to the pronounced hydrophobicity of PET fibers. We therefore hypothesized that a similar behavior could occur with PAs. Surfactants can effectively reduce the PA/water interfacial tension and promote enzyme adsorption, while the addition of co-solvents to the reaction medium may induce polymer swelling and partial solvation of the amide chains, thereby facilitating enzyme accessibility and improving catalytic efficiency. Inspired by previous work on optimizing enzymatic PET and lignocellulose hydrolysis,^[36,44–47]^ we selected a range of commercially available surfactants and co-solvents with potential to enhance Nylon depolymerization, and we evaluated their effect on the hydrolysis titer of Nyl50 using PA66 as substrate. Under our optimized reaction conditions, the addition of 0.5 mM sodium dodecyl sulfate (SDS) or 5 mM 4-dodecylbenzenesulfonic acid both led to approximately 3-fold higher hydrolysis titer compared to control reactions without surfactant. In contrast, the addition of Tween 80 or 4-dodecylphenol (0.5–5 mM) had no measurable impact on product titers (Figure S8). Screening various co-solvents (5–20% v/v) also failed to produce major improvements. However, Nyl50 maintained comparable activity up to ∼10% (v/v) across most of the co-solvents tested (Figure S9A and S9B), except in isopropanol where the enzyme showed low activity even at 5% v/v. A comparison of enzyme stability revealed differences among the homologous nylon hydrolases: Nyl10 retained its activity up to 20% (v/v) DMSO, whereas Nyl50 activity declined sharply beyond 10% (v/v). Cyclopentyl methyl ether (CPME) could also represent a suitable alternative to conventional solvents, as its addition slightly improved the reaction titer up to the highest concentration tested (20%), as shown with Nyl50 and Nyl12 (Figure S9B). Although further studies are needed to rationalize these observations, this represents the first systematic screening of nylon hydrolases under diverse surfactant and co-solvent conditions. These findings provide a useful foundation for future optimization of enzymatic Nylon depolymerization processes. Moreover, the distinct solvent tolerance profiles observed suggest opportunities for hybrid chemo-enzymatic strategies and offer valuable starting points for targeted and rational enzyme engineering efforts.

### Proof of concept of the partial enzymatic degradation on industrially relevant PAs based materials

We next investigated the ability of Nyl50 to hydrolyse industrially-relevant polyamide materials. Most studies on enzymatic hydrolysis of PA have focused on pure or low complexity substrates. However, real polymers are most often contained within complex, multi-material assemblies, films, and/or blends from which isolation of any one material using physical methods alone can be cost prohibitive.^[48]^ In addition, due to their tunability, plastics often contain functional additives/fillers (i.e., thermal- or UV-stabilizers, coatings/paints, fibers/fillers) that can alter or significantly hinder industrial deconstruction and recycling.^[49,50]^ In these contexts, the use of enzymes represents a particularly promising approach, since enzymes evolved for catalytic selectivity even in complex environments.

Here, we evaluated the capacity of Nyl50 to hydrolyze electrical-grade PA66 containing 30% glass fiber and a heat stabilization package. Similar materials are used in automotive and electrical applications such as connectors, bobbins, and polymeric housings. To test the potential application of Nyl50 in the presence of such additives, 10 mg of industrially relevant PA66 or pure PA66 were incubated with pure enzyme (0.3 mg/mL) for 24 h at 75 °C. The soluble fraction of the reaction mixture was recovered and analyzed by IDOT/OPSI-MS. The results showed that Nyl50 exhibited no significant difference between hydrolysis titers on both pure PA66 and industrially relevant PA66 (∼1500 ug/mL), despite the presence of additives or fillers that frequently reduce activity of chemical recycling methods (Figure 4A).^[51]^

**Figure 4.**
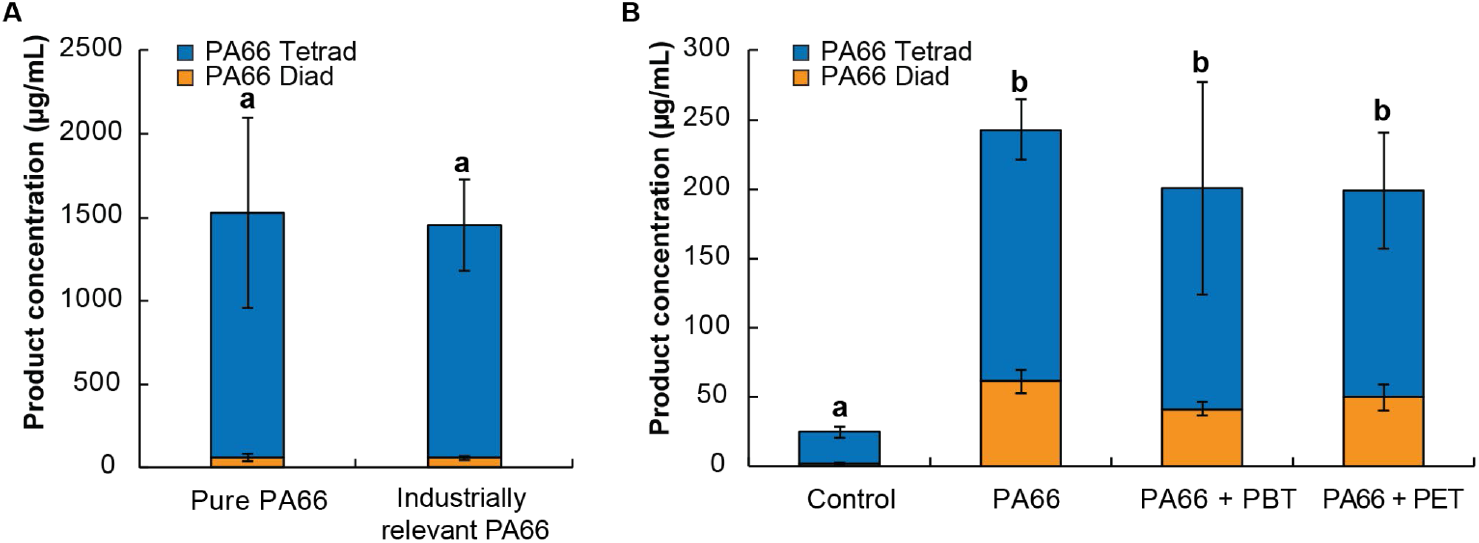
Enzymatic depolymerization reactions of industrially relevant Polyamides with Nyl50. (A) Comparison between pure PA66 and industrially relevant PA66 (B) Effect on a mixture of PA66 and PBT or PET. Reactions were performed with 0.3 mg/mL of pure enzyme in 200 mM Tris pH 8 for 24 h at 75 °C. Error bars show the standard deviation, calculated from two biological replicates. Variability between samples was assessed by pairwise F-tests and differences in means by Welch’s *t*-tests. Samples sharing the same letters indicate no significant difference (*p* > 0.05).

Next, given that the industrially relevant polyamide material is often embedded within a matrix of other polymers such as polybutylene terephthalate (PBT), we aimed to evaluate the activity of Nyl50 for hydrolysis of a composite substrate containing PA66 and PBT. To better approximate realistic conditions for polymer degradation, enzymatic assays were conducted using a 50:50 (w/w) mixture (5 mg each) of PA66 with either PBT or PET. Here again, PA66 hydrolysis titers remained consistent, regardless of the presence of PBT or PET (Figure 4B). Similarly, although not the primary focus of this study, the activity of PETase (LCC wt, single mutant 207 and single mutant 192)^[52]^ on PET and PBT was also assessed in the presence of PA66, under the same conditions and analyzed via colorimetric assay. The observed titers were likewise unaffected by the addition of 50% PA66 (Figure S10).

Taken together, these results indicate that Nyl50 retains its catalytic efficiency in the presence of complex polymer mixtures and highlight its potential applicability in the enzymatic recycling of mixed polyamide-based materials. Although the degree of depolymerization remains far too low to be viable or implemented at a larger scale, the combination of an effective pretreatment and future enzyme engineering campaigns could help move closer to that goal.

## Conclusion

In this work, we optimized the reaction conditions for three recently discovered nylon hydrolases. While all three enzymes demonstrated significant improvements in activity using optimized conditions, hydrolytic titers using Nyl12 improved to 4 % and 2 % depolymerization on PA6 and PA66, respectively. Nyl12 thus outperforms all enzymes currently described in the literature, including those reported on pretreated substrates or involving engineered enzymes. We also demonstrated that enzymatic Nylon hydrolysis is unaffected by the presence of additives, fillers, and copolymers typically used in automotive and electrical applications. Although the overall degree of depolymerization remains modest, we demonstrate that substrate modification, specifically by adjusting particle size and surface area, can significantly enhance enzymatic activity, as well as by the addition of surfactants. As highlighted in analogous studies on PET, the development of efficient pretreatment strategies appears to be critical for efficient enzymatic Nylon recycling. Finally, this work provides a valuable foundation for the development of future depolymerization processes and future enzyme engineering campaigns.

## Materials and Methods

### Substrate preparation

PA6 and PA66 pellets were processed into fine powders via cryomilling at −196 °C in a Retsch CryoMill using a stainless steel jar and steel grinding media. Approximately 2 to 10 grams of pellets with a 25 mm grinding ball were loaded into the jar and pre-cooled at 5 Hz for approximately 5 minutes under liquid nitrogen. After equilibration at −196 °C, the jar was then shaken at 30 Hz for 30 minutes, rested for 5 minutes under a liquid nitrogen flow, and shaken again at 30 Hz for 30 minutes. To obtain finer powders, the procedure was repeated using 5-6 mm grinding balls instead of the 25 mm ball. After warming to near room temperature, the jar was opened and the powders were recovered. The ball-milled powders were sieved through 450, 230, and 140 mesh sieves (VWR) using deionized (DI) water to aid passage through the sieves. The collected powders were subsequently freeze-dried to remove residual moisture. To eliminate unreacted precursors and byproducts, the dried powders were washed three times with methanol following a reported procedure.^[43]^ Finally, the washed powders were vacuum-dried and used as obtained for further experiments.

### Enzyme expression & purification

Enzymes were expressed and purified following the previous reported methods,^[19]^ with no modifications. Expression of LCC wt, LCC 2017 and LCC 192 was done with the same procedure.

### Optimization of the reaction conditions

Reactions to determine the optimal temperature were performed in 20 mM Phosphate, pH 7.4, for 24 h and 48 h. The temperatures tested were 35 °C, 45 °C, 55 °C, 65 °C, 75 °C and 85 °C. Reactions to determine the optimal buffer and pH were performed at 65 °C using acetate at pH 4 or pH 5; Phosphate at pH 6, pH 7, or pH 8; and Tris at pH 6, pH 7, pH 8, or pH 9. All pHs were measured at room temperature. Reactions were carried out with 10 mg of PA in 75 uL buffer and 25 ul enzyme lysate, for 12 h or 24 h. Each enzymatic reaction and parameter were run in triplicate, treated separately and compared with a reference setup (65°C, 20 mM phosphate buffer, pH 7.4), referred to as “original” conditions.

### In vitro Enzyme Activity Assays under optimal conditions

Reactions were carried in triplicate at 100 μL, containing 20 mM Phosphate (pH 7.4) or 200 mM Tris (pH 8), 0.3 mg/mL of pure enzyme, and 10 mg of PA66 or PA6 as powder form and was incubated at 65 °C or 75 °C for 12 h to 72 h without shaking. The samples were centrifuged at 4,500 rpm for 10 min, and the supernatant was subjected to I.DOT/OPSI-MS analysis to determine the formation of corresponding hydrolysis products. The degree of depolymerization was calculated in accordance with our previous study.^[19]^

### Scale-up

Reaction scale-up was carried out using a constant concentration of substrates (10 mg/100 uL) and enzymes, using the same lysate and similar reaction conditions. Initial reactions were run, in triplicate, in a 100 μL volume reaction, containing 200 mM Tris (pH 8), 25 μL of heat-treated lysate, and 10 mg of PA66 as powder form and were incubated in at 75 °C for 4 h in a PCR tube without shaking. Reactions were then scaled up in a 2 mL Eppendorf tube, in triplicate, with 100 mg of PA66 powder in 1 mL final volume. Finally, one reaction was run in a 50 mL round bottom flask with constant stirring (250 rpm), containing 1 g of PA66 powder in a 10 mL final volume. All samples were centrifuged at 4,500 rpm for 10 min, and the supernatant was subjected to I.DOT/OPSI-MS analysis to determine the formation of corresponding hydrolysis products.

### Metal ions, surfactants and solvents screening

Reactions to determine the influence of metal ions were carried out in triplicate in a total volume of 100 μL, containing 200 mM Tris buffer, pH 8 (optimal) and 20 mM Phosphate buffer, pH 7.4 (original), 25 μL of enzyme lysate, and 10 mg of substrate powder.

For the surfactants and solvents screening, reactions were carried out in triplicate in a total volume of 100 μL, containing 200 mM Tris buffer (pH 8), 0.3 mg/mL purified enzyme or 25 μL of enzyme lysate, and 10 mg of substrate powder. The mixtures were incubated at 75 °C for 6 h without agitation in the presence of either 0.5, 1, or 5 mM of sodium dodecyl sulfate (SDS), Tween 80, 4-dodecylphenol, 4-dodecylbenzenesulfonic acid, or lauric acid, or with 5, 10, 15, or 20 % (v/v) of dimethyl sulfoxide (DMSO), methanol (MeOH), acetone, dimethylformamide (DMF), isopropanol, or cyclopentyl methyl ether (CPME). Following incubation, samples were centrifuged at 4,500 rpm for 10 min, and the supernatants were analyzed by mass spectrometry (MS) to quantify the formation of corresponding hydrolysis products. Results were compared to control reactions performed under identical conditions without any surfactant or co-solvent. Negative controls without enzyme were also included for each tested condition. All surfactants and co-solvents were purchased from Sigma-Aldrich.

### In vitro Enzyme Activity Assays with industrially relevant polyamides

A typical enzymatic reaction was performed in duplicate at a final volume of 100 μL containing 200 mM Tris (pH 8), 25 uL 0.3 mg/mL of pure enzyme, and 10 mg of PA66 or 5 mg of PA66 mixed with 5 mg of PBT or PET (SigmaAldrich). Reactions were incubated at 75 °C for 12 h without shaking. The samples were centrifuged at 4,500 rpm for 10 min, and the supernatant was subjected to I.DOT/OPSI-MS to determine the formation of corresponding hydrolysis products. Statistical analysis of variance was performed using pairwise F-tests to compare variability between sample groups, and Welch’s *t*-tests were applied for mean comparisons (both two-tailed, assuming unequal variances), following standard procedures for small-sample analysis. ^[53]^

### I.DOT/OPSI-MS analysis of enzyme activity

The immediate drop-on-demand technology (Dispendix GmbH, Stuttgart, Germany) coupled with open port sampling interface mass spectrometry (I.DOT/OPSI-MS) was used to analyze enzymatic reactions of nylons with high throughput as previously described in detail.^[54]^ Enzymatic reactions were diluted 1:100 (v/v) in HPLC grade water containing 500 nM propranolol which served as an internal standard for droplet detection and droplet normalization. 40 µL of diluted reactions were transferred to I.DOT S.100 96-well plates and analyzed immediately by ejecting 20 nL of sample into the OPSI-MS having a flow of 75/25/0.1 (v/v/v) acetonitrile/water/formic acid. Samples were transported to the electrospray ion source of a Sciex 7500 triple quadrupole mass spectrometer (SCIEX, Concord, Ontario, Canada) for ionization and detection. In-house developed softwares (ORNL I.DOT-MS Coupler v2.50) were used for control of the IDOT system, extraction of data from vendor file formats, peak finding, and peak integration. Each droplet signal was background subtracted and normalized to propranolol signal. Absolute quantitation was achieved using a 12-point calibration curve for each oligomer (0-50 µg/mL). Each sample was measured in triplicate, with the means averaged across sample replicates for reported means and standard deviations. The 7500 was operated in positive ion mode with flow = 170 µL/min, gas setting 1 = 90, gas setting 2 = 60, electrospray voltage = 5.5 kV, capillary temperature = 200 °C, and dwell time = 20 ms. Multiple reaction monitoring (MRM) used the following transitions: 260.1→183.0; collision energy (CE)=26 eV (propranolol), 245.19→100.11; CE=27 eV (PA66-linear monomer, diad), 471.35→100.11; CE=65 eV (PA66-linear dimer, tetrad), 245.19→114.09; CE=26 eV (PA6-linear dimer), 358.27→114.09; CE=50 eV (PA6-linear trimer), and 471.35→114.09; CE=65 eV (PA6-linear tetramer). Measurements of PA66 and PA6 oligomers were always collected in separate experiments.

### Brunaner-Emmett-Teller (BET) surface area measurement of ball-milled PA66 powder

Nitrogen adsorption isotherms were measured using a Micromeritics 3Flex Adsorption Analyzer, and the data were processed with the accompanying software (3Flex Version 5.03). All samples for nitrogen isotherm measurements were loaded into a tared sample tube set (tube and check valve) and degassed at 50 °C overnight on a Micromeritics VacPrep system. BET surface areas were determined from multipoint linear regression within the relative pressure p/p° range of 0.01-0.9, yielding correlation coefficients greater than 0.999.

## Contributions

C.B: Conceptualization; biochemistry; writing. L.X.Z: Biochemistry. D.L.C: Analytical characterization. R.L.C: Ball-milling of the nylon powder. M.Z, V.B: Substrate preparation (washing and sieving) and characterization; writing. L.W., JHBH: Analysis of industrially-relevant samples. M.K: Project coordination; writing editing. J.F.C: Analytical methods development and analysis; supervision of analytical research. J.K.M: Conceptualization; funding acquisition; supervision, writing.

