## Supplemental Information for "Process optimization for enhanced enzymatic nylon deconstruction"

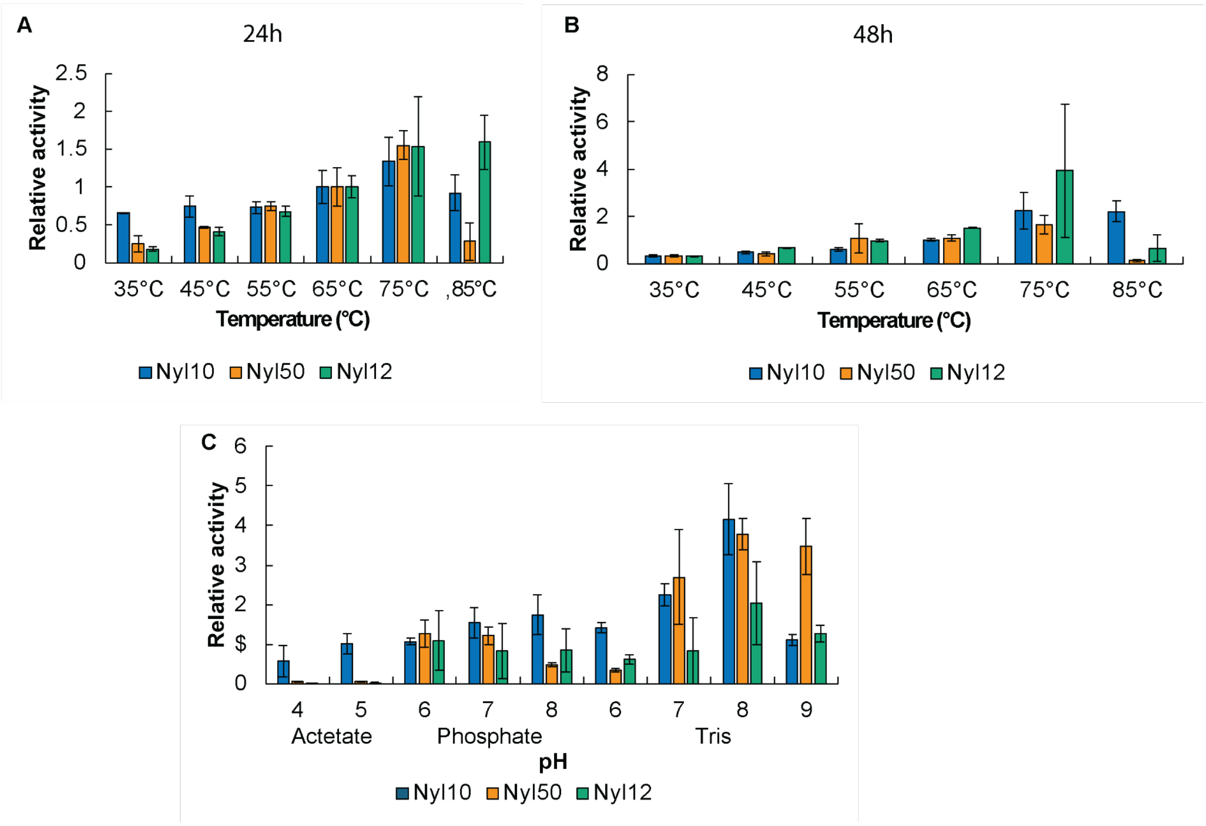


**Figure S1. Enzymatic PA66 depolymerization reactions with Nyl10, Nyl50 and Nyl12 in function of temperature and pH.** (A) Effect of temperature on enzymes hydrolysis activity after 24 h. Reactions ran with crude lysate in 20 mM phosphate pH 7.4 for 24 h. (B) Effect of temperature on enzymes hydrolysis activity after 48 h. Reactions ran with crude lysate in 20 mM phosphate pH 7.4 for 48 h. (C) Effect of pH on hydrolysis activity. Reactions ran with crude lysate for 12 h at 65°C. Error bars show the standard deviation, calculated from three biological replicates.

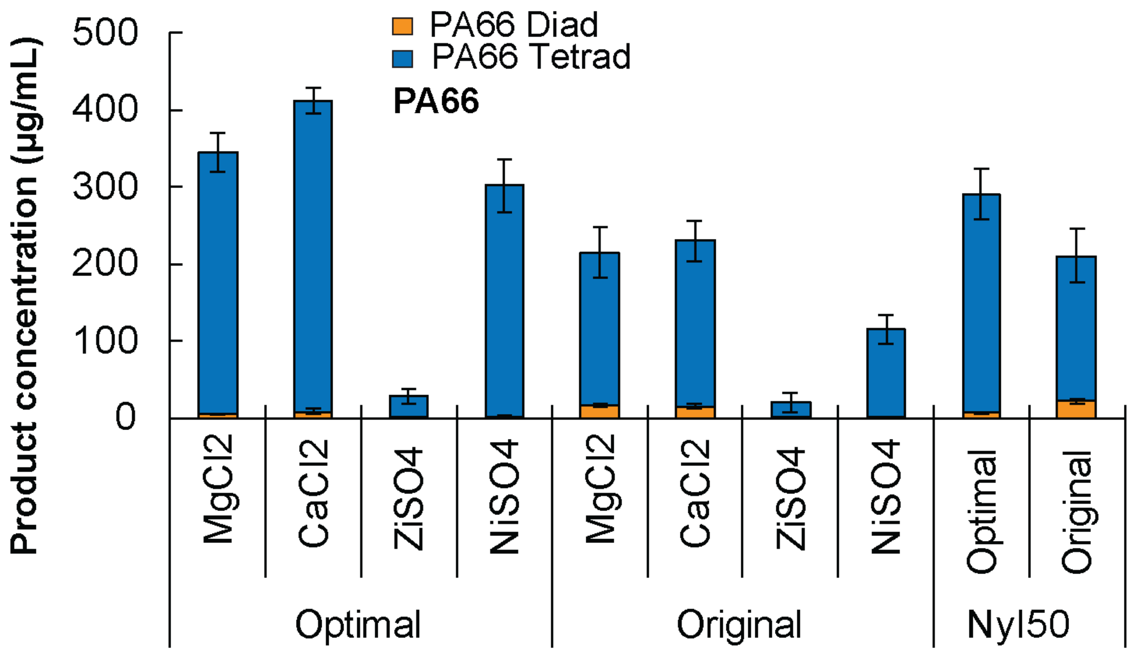


**Figure S2. Influence of metal ions on PA66 depolymerization reactions with Nyl50.** Reactions ran with Nyl50 crude lysate for 24 h at 65°C, with 10 mM $MgCl_2$, $CaCl_2$, $ZnSO_4$, or $NiSO_4$. Error bars show the standard deviation, calculated from three biological replicates.

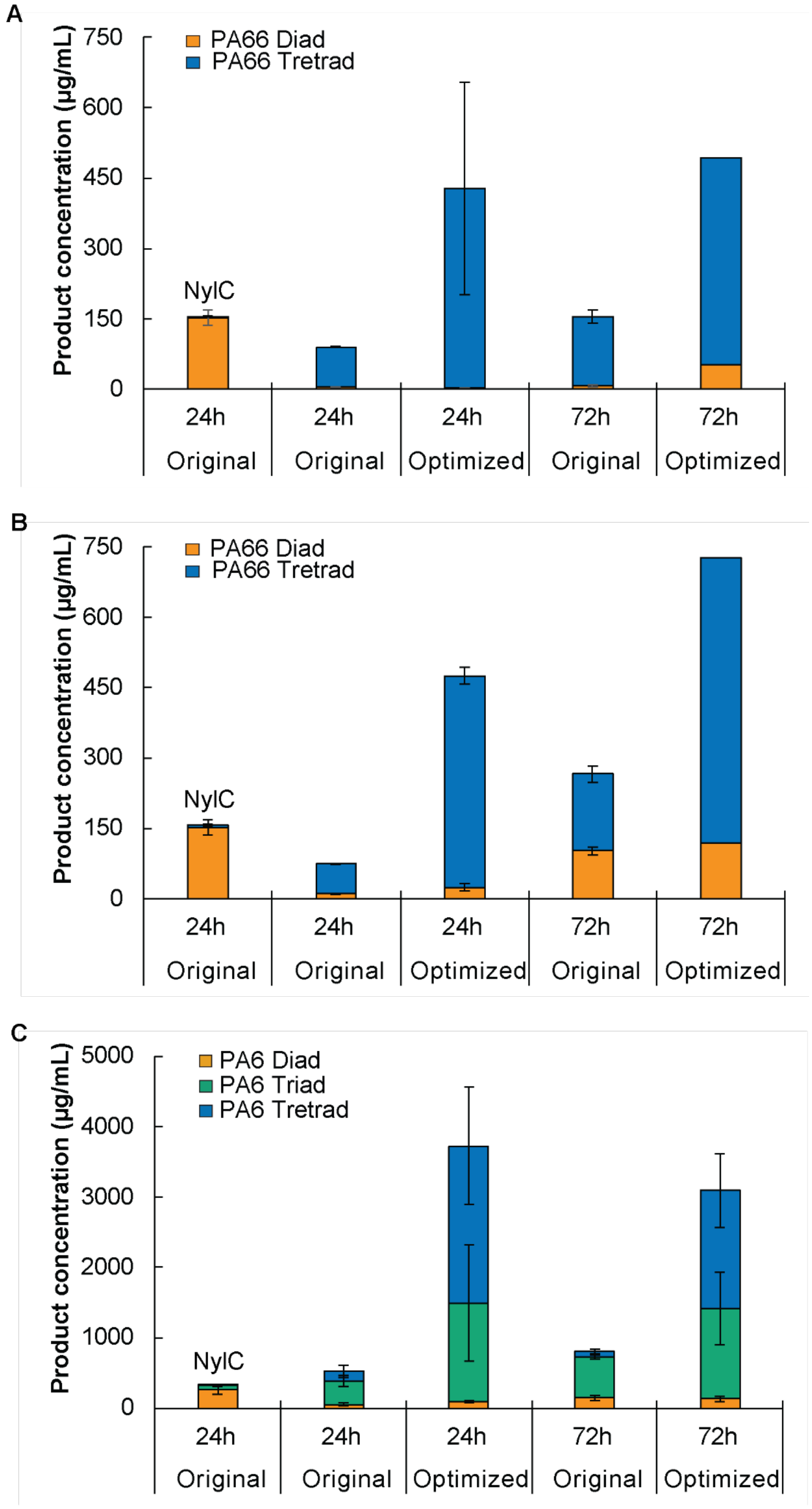


**Figure S3. Enzymatic PA66 and PA6 depolymerization reactions with Nyl50, Nyl10 and Nyl12 under optimal conditions, at 0.3 mg/mL of enzymes.** (A) Nyl50 with PA66 (B) Nyl10 with PA66 (C) Nyl12 with PA6. Reactions ran with 0.3 mg/mL or 1 mg/mL of pure enzyme in 20 mM Phosphate pH 7.4 for 72 h (original) or in 200 mM Tris pH 8 for 24h and 72 h (optimal). Error bars show the standard deviation, calculated from three biological replicates.

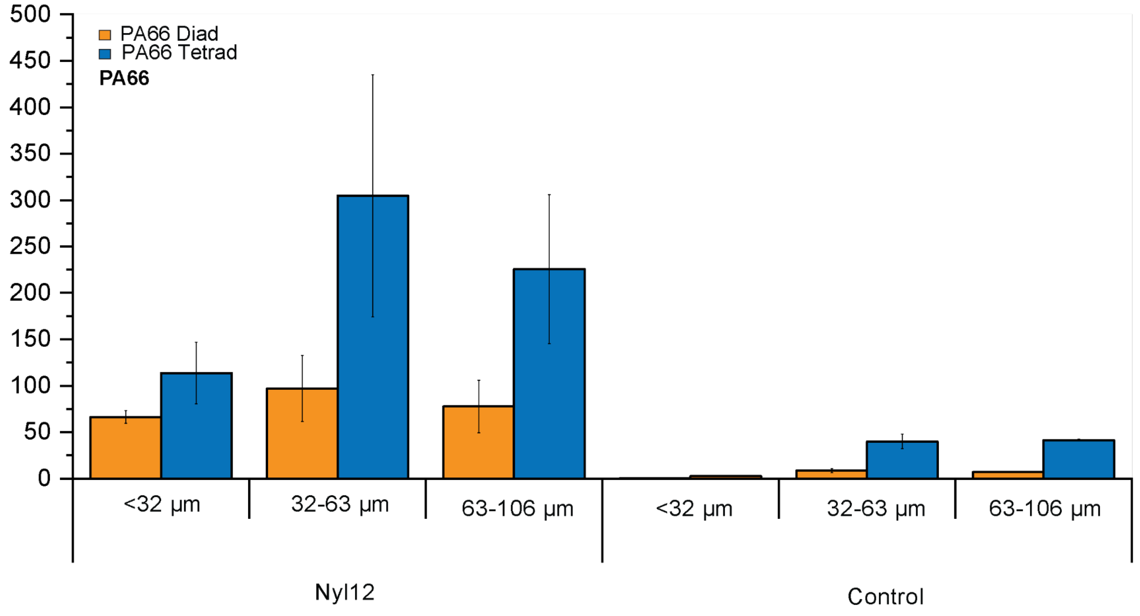


**Figure S4. Determination of substrate properties influencing enzymatic hydrolysis.** Concentration of hydrolysis products *vs.* PA66 sizes in the presence and absence (control) of Nyl12. Reactions ran with crude lysate, 6 h at 75°C. Error bars show the standard deviation, calculated from two biological replicates.

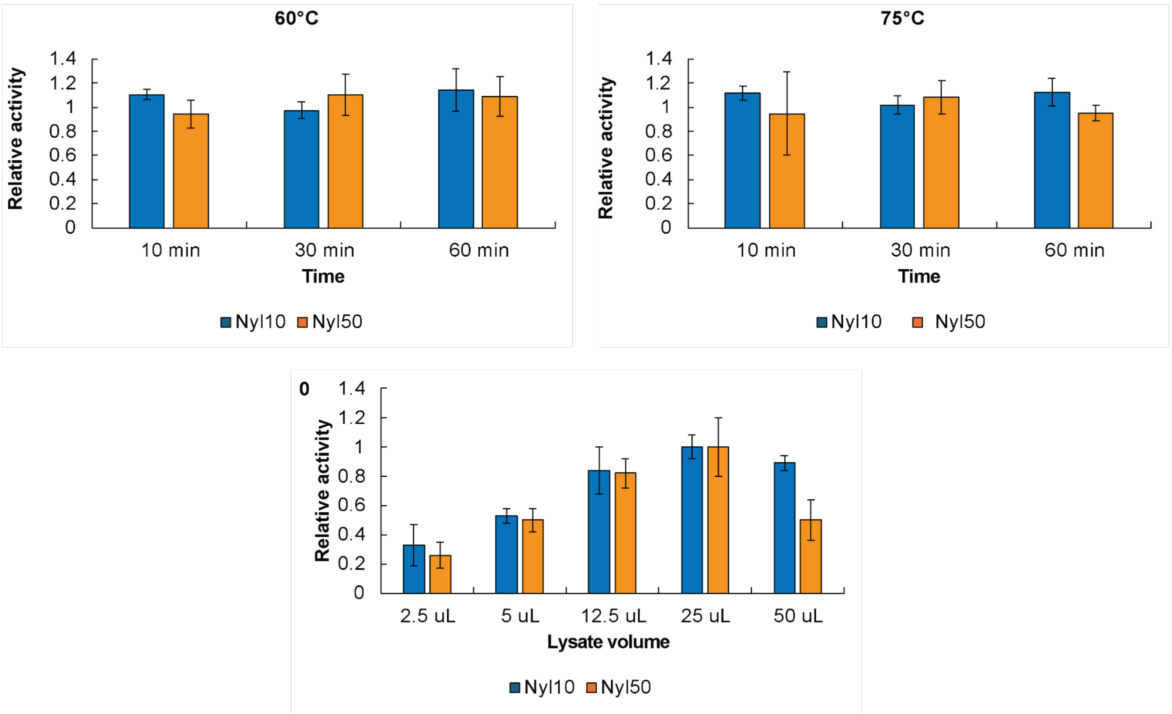


**Figure S5. Effect on PA66 hydrolysis of heat treatment and ratio lysate/buffer concentration with Nyl50 and Nyl10.** Reactions ran with crude lysate. Error bars show the standard deviation, calculated from three biological replicates.

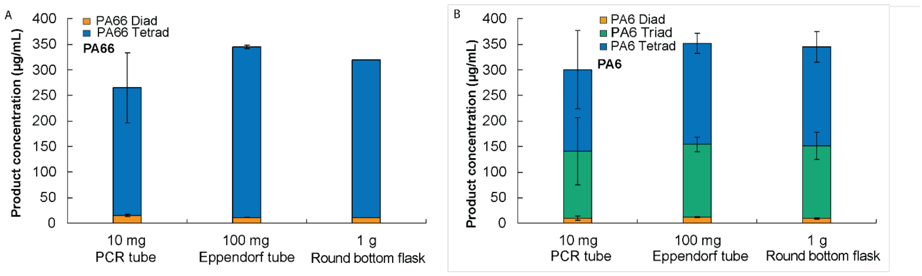


**Figure S6. Enzymatic PA66 reactions scale up with Nyl50 under optimal conditions.** Reactions ran with a heat-treated enzyme lysate in 200 mM Tris pH 8 for 4 h at 75°C. Error bars show the standard deviation, calculated from three biological replicates.

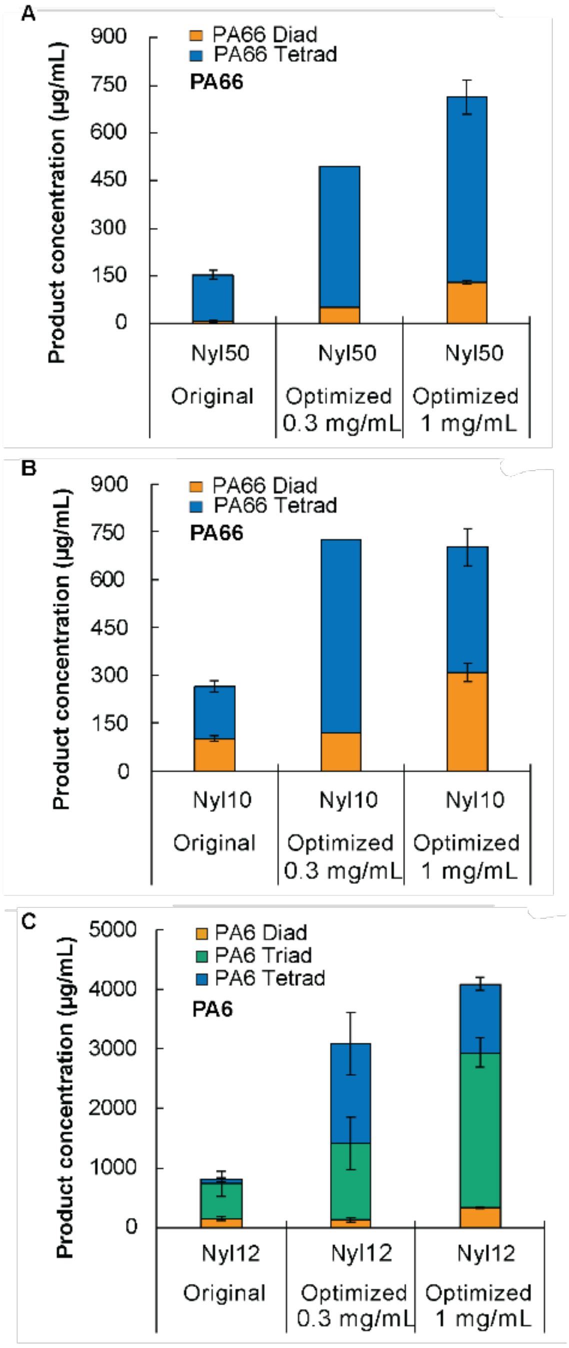


**Figure S7. Enzymatic PA66 and PA6 depolymerization reactions with Nyl50, Nyl10 and Nyl12 under optimal conditions, at 1 mg/mL of enzymes.** (A) Nyl50 with PA66 (B) Nyl10 with PA66 (C) Nyl12 with PA6. Reactions ran with 0.3 mg/mL or 1 mg/mL of pure enzyme in 20 mM Phosphate pH 7.4 for 72 h (original) or in 200 mM Tris pH 8 for 72 h (optimal). Error bars show the standard deviation, calculated from three biological replicates.

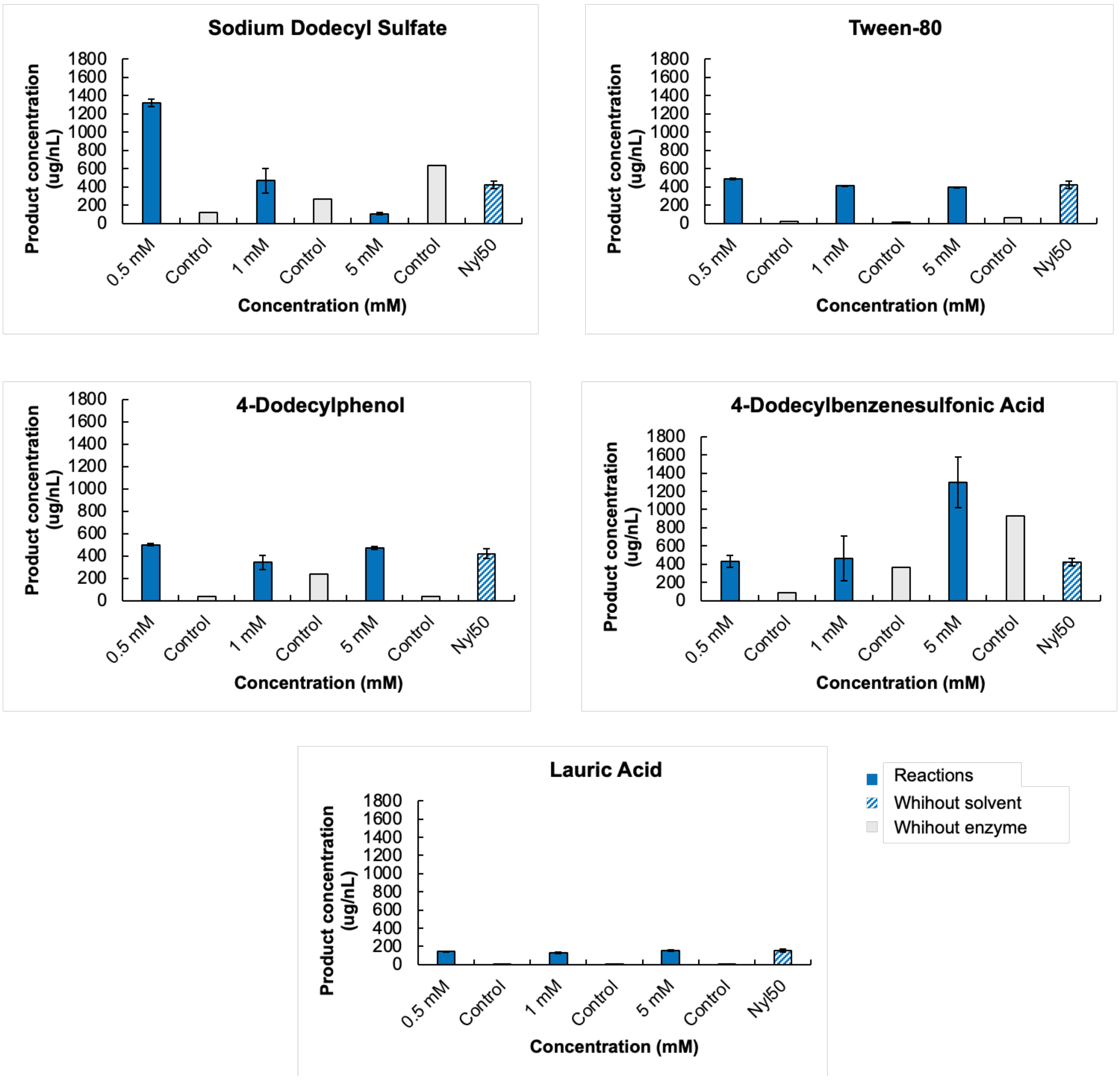


**Figure S8. Influence of surfactants on PA66 depolymerization reactions with Nyl50.** Reactions ran with crude lysate for 6 h at 65°C, in 200 mM Tris buffer, pH 8. In both panels, error bars show the standard deviation, calculated from three biological replicates. Controls are reactions run without enzyme. Nyl50 is a control run under the same conditions without solvents.

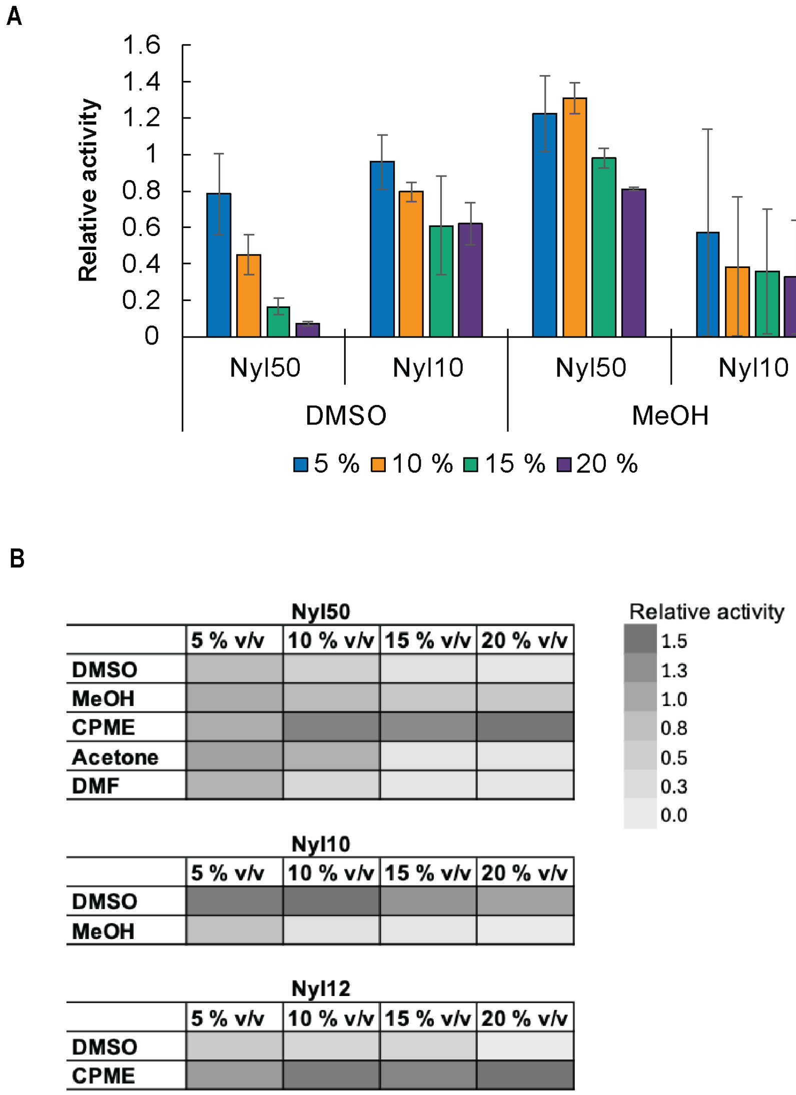


**Figure S9. Influence of solvents on PA66 depolymerization reactions with Nyl50.** Reactions ran with crude lysate for 6 h at 65°C, in 200 mM Tris buffer, pH 8. (A) Comparison between Nyl50 and Nyl10 with DMSO and MeOH. Error bars show the standard deviation, calculated from three biological replicates. (B) Heatmap of the solvents screening for Nyl50, Nyl10 and Nyl12. Control reactions run without enzyme did not show any activity.

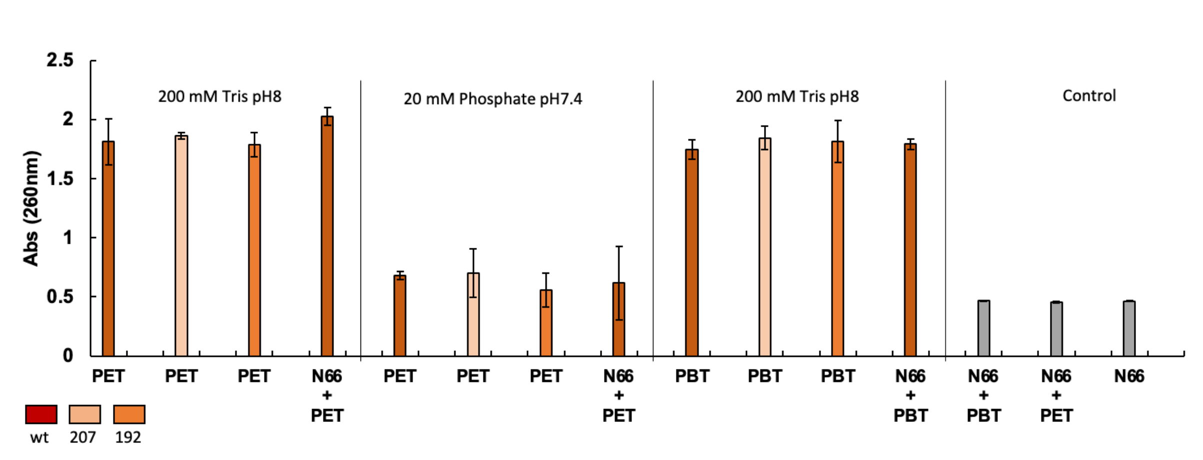


**Figure S10. Enzymatic depolymerization reactions of PET and PBT with ICC variants.** Effect on a mixte of PA66 and PBT or PET. Reactions ran with crude lysate of LCC wt, LCC 207 or LCC 192 mutants in presence or not of PA66, in 20 mM Phosphate pH 7.4 for 72 h (original) or in 200 mM Tris pH 8 for 24h and 72 h (optimal). Error bars show the standard deviation, calculated from two biological replicates.
